# TGF-β Signaling in Cranial Neural Crest Affects Late-Stage Mandibular Bone Resorption and Length

**DOI:** 10.1101/2024.05.24.595783

**Authors:** Claire J. Houchen, Saif Ghanem, Vesa Kaartinen, Erin Ealba Bumann

## Abstract

Malocclusions are common craniofacial malformations which cause quality of life and health problems if left untreated. Unfortunately, the current treatment for severe skeletal malocclusion is invasive surgery. Developing improved therapeutic options requires a deeper understanding of the cellular mechanisms responsible for determining jaw bone length. We have recently shown that neural crest mesenchyme (NCM) can alter jaw length by controlling recruitment and function of mesoderm-derived osteoclasts. Transforming growth factor beta (TGF-β) signaling is critical to craniofacial development by directing bone resorption and formation, and heterozygous mutations in TGF-β type I receptor (*TGFBR1)* are associated with micrognathia in humans. To identify what role TGF-β signaling in NCM plays in controlling osteoclasts during mandibular development, mandibles of mouse embryos deficient in the gene encoding *Tgfbr1* specifically in NCM were analyzed. Our lab and others have demonstrated that *Tgfbr1*^*fl/fl*^*;Wnt1-Cre* mice display significantly shorter mandibles with no condylar, coronoid, or angular processes. We hypothesize that TGF-β signaling in NCM can also direct later bone remodeling and further regulate late embryonic jaw bone length. Interestingly, analysis of mandibular bone through micro-computed tomography and Masson’s trichrome revealed no significant difference in bone quality between the *Tgfbr1*^*fl/fl*^*;Wnt1-Cre* mice and controls, as measured by bone perimeter/bone area, trabecular rod-like diameter, number and separation, and gene expression of Collagen type 1 alpha 1 (*Col1α1*) and Matrix metalloproteinase 13 (*Mmp13*). Though there was not a difference in localization of bone resorption within the mandible indicated by TRAP staining, *Tgfbr1*^*fl/fl*^*;Wnt1-Cre* mice had approximately three-fold less osteoclast number and perimeter than controls. Gene expression of receptor activator of nuclear factor kappa-β (*Rank*) and *Mmp9*, markers of osteoclasts and their activity, also showed a three-fold decrease in *Tgfbr1*^*fl/fl*^*;Wnt1-Cre* mandibles. Evaluation of osteoblast-to-osteoclast signaling revealed no significant difference between *Tgfbr1*^*fl/fl*^*;Wnt1-Cre* mandibles and controls, leaving the specific mechanism unresolved. Finally, pharmacological inhibition of *Tgfbr1* signaling during the initiation of bone mineralization and resorption significantly shortened jaw length in embryos. We conclude that TGF-β signaling in NCM decreases mesoderm-derived osteoclast number, that TGF-β signaling in NCM impacts jaw length late in development, and that this osteoblast-to-osteoclast communication may be occurring through an undescribed mechanism.

## 1 Introduction

Craniofacial anomalies are a diverse group of malformations in the growth of skull and facial bones. These anomalies can be inherited or spontaneous, and both genetic and environmental factors have important influences on craniofacial development (1). Craniofacial anomalies present a significant clinical issue because the only treatment option currently available is invasive surgery (2, 3). The jaw skeleton, specifically, displays a wide range of anomalies including hypo- and hyperplasia, pro- and retrognathia, and micro- and macroganthia. Increasing our understanding of the molecular and cellular mechanisms by which elements of the jaw skeleton achieve their appropriate shape and size also increases our ability to develop minimally invasive therapeutic options to improve quality of life for patients with jaw anomalies.

During embryonic development, neural crest mesenchyme (NCM) forms all the bony elements in the facial and jaw skeletons. NCM-derived osteoblast lineage cells form the bone and NCM-derived chondrocyte lineage cells form the cartilage of the facial and jaw bones (4). NCM is known to specifically be involved in osteogenic induction, proliferation, differentiation, mineralization, and matrix remodeling (5-8). It is well-established that osteoblast lineage cells such as osteocytes influence osteoclastogenesis and osteoclast function, and there is growing evidence that bone resorption by osteoclasts is critical to mandibular development (9-11). It is therefore logical that in addition to its long-known role in facial and jaw osteogenesis, NCM also regulates jaw length by controlling bone resorption, which has been demonstrated in both mice and birds (5, 12, 13). The primary cell type known to play a role in bone resorption is mesoderm-derived osteoclasts, thus not an NCM-derived cell. An understanding of embryonic cellular signaling, particularly between NCM-derived osteoblast lineage cells and mesoderm-derived osteoclasts, is critical for understanding the origin of jaw bone anomalies in human patients.

One pathway which may be involved in communication between NCM-derived osteoblast lineage cells and mesoderm-derived osteoclasts is TGF-β signaling. The TGF-β signaling pathway is comprised of a heteromeric complex of two type one and two type two transmembrane serine/threonine kinases, TGFBR1 and TGFBR2 (14). Upon ligand binding, the activated TGF-β receptor complex phosphorylates Smad proteins that form a signal cascade which accumulates in the nucleus, where they serve as transcription factors (15, 16). The TGF-β signaling family is known to play an important role in embryonic craniofacial development, cell proliferation, migration and differentiation, and extracellular matrix secretion (17-22). Within bone tissue, TGF-β is known to regulate many aspects of bone remodeling, such as osteoblast proliferation, osteoblast differentiation, and osteoclast apoptosis, through modulation of factors such as matrix metalloproteinase 13 (19, 23). Consequently, patients who are heterozygous for mutations in *TGFBR1* have Loeys-Dietz Syndrome I, which is associated with craniofacial changes such as craniosynostosis, facial asymmetry, low set ears, hypertelorism, cleft palate, and microretrognathia (24-26). Therefore, TGF-β signaling in NCM is critical during facial and jaw development, but it is not yet fully understood.

Previous studies have begun to explore TGF-β signaling in NCM and jaw development, and specifically, both TGFBR1 and TGFBR2 have emerged as regulators of mandible patterning (20-22). Loss of *Tgfbr1* in NCM results in a shortened mandible and a lack of development of the coronoid, condylar, and angular processes (17, 27). Previously, the mandibular defect in mice lacking *Tgfbr1* in NCM was attributed to high rates of apoptosis from stage E10 to E14.5 in the proximal and aboral regions of the mandible, while no changes in cell proliferation were noted (17, 27). In this study, we elucidate the mechanism of NCM control of jaw development by modulating TGF-β signaling in NCM and investigating resulting bone resorption changes in the lower jaw. We hypothesized that in addition to high rates of apoptosis in the jaws of mice that lack *Tgfbr1* in NCM, osteoclastogenesis and bone resorption would be altered in the mandible due to loss of typical TGF-β signaling in NCM-derived osteoblasts and osteocytes.

## 2 Materials and Methods

### 2.1 Generation and genotyping of *Tgfbr1*^*fl/fl*^*;Wnt1-Cre* mice

*Alk5*^*fl/fl*^ female mice (28) were mated with *Wnt1-Cre* male mice (29) to generate the *Alk5*^*fl/+*^*/Wnt1-Cre* mice, and then the *Alk5*^*fl/fl*^ female mice were mated with *Alk5*^*fl/+*^*/Wnt1-Cre* male mice to generate the experimental *Tgfbr1*^*fl/fl*^*;Wnt1-Cre* embryos (17, 27). *Tgfbr*^*fl/+*^ mice were used as experimental controls. All mice used in these studies are on a mixed C57BL6/J 129Sv background.

Genotyping was done using a MultiGene™ Gradient PCR Thermal Cycler (Labnet, Edison, NJ) or Veriti™ 96-Well Fast Thermal Cycler (ThermoFisher Scientific, Wilmington, DE) on tail tip and yolk sac DNA. Mutant embryos were identified by PCR genotyping for presence of the *cre* transgene (Forward 5’-CGTTTTCTGAGCATACCTGGA-3’ and Reverse 5’-ATTCTCCCACCGTCAGTAGG-3’) and the *Tgfbr1* floxed alleles (Forward 5’-GAGTCTGAAGCTTTGCAAG-3’ and Reverse 5’-ATTAGCTAAGCCCCTTC-3’).

For staging, the noon on the plugging day was designated as E0.5. Embryos were staged by comparison with references (30). Mouse embryos were collected for analysis at stages E16.5 and E18.5. All protocols were approved by the Institutional Animal Care and Use Committee at the University of Michigan (PRO00008469).

### 2.2 Microcomputed tomography analysis of mandibular bone

Specimens placed in a 19 mm diameter specimen holder and scanned over the entire skull using a microcomputed tomography (micro-CT) system (µCT100 Scanco Medical, Bassersdorf, Switzerland). The following scan settings were used: voxel size 10 µm, 55 kVp, 109 µA, 0.5 mm AL filter, and integration time 500 ms. Analysis of bone volume and tissue mineral density was performed using the manufacturer’s evaluation software. A fixed global threshold of 13% (130 on a grayscale of 0–1000) for E16.5 mice and 15% (150 on a grayscale of 0-1000) for E18.5 mice was used to segment bone from non-bone. Four control and four *Tgfbr1*^*fl/fl*^*;Wnt1-Cre* samples were assessed at both E16.5 and E18.5 (n=4/genotype/stage).

### 2.3 Whole-mount staining of tartrate-resistant acid phosphatase

To detect bone resorption, embryos were fixed in 10% formalin overnight at 4°C and stained to identify tartrate-resistant acid phosphatase (TRAP) activity. Samples were stained whole-mount using an Acid Phosphatase Leukocyte Kit following the manufacturer’s protocol, except Fast Red Violet was used in place of Fast Garnet GBC Base Solution (Sigma-Aldrich, Saint Louis, MO) (5). Seventeen control and nine *Tgfbr1*^*fl/fl*^*;Wnt1-Cre* samples were assessed from four litters. Images of the embryos were taken on a MZ95 microscope (Leica, Buffalo Grove, IL) with a DP72 camera using cell Sens Entry software (Olympus, Center Valley, PA).

### 2.4 Histology

To examine bone quality and bone resorption on the cellular level, embryos were fixed in 10% formalin overnight at 4°C, washed in PBS, decalcified in EDTA/Tris Buffered Saline for 1-3 days at 4°C, dehydrated, embedded in paraffin, and cut into 7 µm transverse sections. To analyze mandibular bone quality, sections were stained with Masson’s trichrome: Bouin’s fluid overnight, Mayer’s Hematoxylin for 10 min, Biebrich scarlet for 2 min, phosphomolybdic acid/phosphotungstic acid for 15 min, and aniline blue for 15 min. To identify bone resorption in the mandible, adjacent sections to each Masson’s trichrome were stained with TRAP as described above for the Whole-mount embryos and counterstained with fast green (Vector Laboratories, Burlingame, CA).

Images were obtained on a BX51 upright light microscope with a DP70 camera using DP Controller software (Olympus) or an Axiovert 200M microscope with an AxioCam MRx using AxioVision 4.8 software (Zeiss, Thornwood, NY). For the images collected with the DP controller software, Image Composite Editor software (Microsoft Research, Redmond, WA) was used to stitch multiple images into one.

Bioquant Osteo 2014 v141.60 software (Bioquant Image Analysis Corporation, Nashville, TN) was used to quantify mandibular bone quality using Masson’s trichrome-stained sections and to quantify mandibular bone resorption using TRAP/fast green-stained sections. Bone quality was assessed by the following measures: bone perimeter per bone area (B.Pm/B.Ar), trabecular rod-like width (Tb.W), trabecular number (Tb.N), and trabecular separation (Tb.Sp). Bone resorption was assessed by osteoclast perimeter per bone area (Oc.Pm/B.Ar) and number of osteoclasts per bone area (Oc.N/B.Ar) in sections adjacent to sections used for bone quality analysis (31). Three control and three *Tgfbr1*^*fl/fl*^*;Wnt1-Cre* E18.5 embryos from the same litter were assessed.

### 2.5 Gene expression analysis by RT-qPCR

Expression of genes related to osteoblasts, osteoclasts, and osteoblast-to-osteoclast signaling was assessed by reverse transcription quantitative PCR (RT-qPCR) (32). Total RNA was isolated from E18.5 mandibles with the soft tissue removed using an RNeasy column purification kit (Qiagen, Valencia, CA). The ascending ramus was removed in control samples because *Tgfbr1*^*fl/fl*^*;Wnt1-Cre* lack the ascending ramus and all mandibular processes (33-36). Concentration and purity of RNA were assessed using a Nanodrop ND-1000 (ThermoFisher Scientific). Approximately 560ng of total RNA was converted to cDNA in a 20μl reverse transcription reaction using the Omniscript RT kit (Qiagen) and random primers (ThermoFisher Scientific). The reaction was processed for 90 minutes at 37°C and then stored at -20°C.

RT-qPCR was performed in a ViiA™ 7 Real-Time PCR System (ThermoFisher Scientific, Grand Island, NY). Forward and reverse primers, 1 μl of cDNA, RNase-free dH20, and TaqMan Universal PCR Master Mix (ThermoFisher Scientific), containing dNTPs with dUTP, AmpliTaq Gold DNA polymerase, Uracil-DNA Glycosylase, Passive Reference 1 and optimized buffer components, were manually mixed in a 30 μl reaction to amplify the cDNA of interest. Samples were run on MicroAmp Optical 96-well Reaction plates (ThermoFisher Scientific). The thermal cycling parameters for all primers was as follows: step 1, 50°C for 2 minutes; step 2, 95°C for 10 minutes; step 3, 60°C for 1 minute and a plate read; step 3 was repeated 40 times; step 4, 60°C for 1 minute. PCR products amplified after 35 cycles were considered to be false positives. Roche Applied Science ProbeFinder was used to design the following mouse gene primers: *β-actin* (Forward 5’– TGACAGGATGCAGAAGGAGA–3’ and Reverse 5’–CGCTCAGGAGGAGCAATG–3’), used with Universal primer #106 with an amplicon length of 75 bp and Collagen type 1 alpha 1 (*Col1□1*) (Forward 5’–CATGTTCAGCTTTGTGGACCT–3’ and Reverse 5’– GCAGCTGACTTCAGGGATGT–3’), used with Universal primer #15, with an amplicon length of 94 bp. Primers for mouse *Mmp13, Mmp9, Rank, Rankl, Opg*, and *M-csf* are commercially available from ThermoFisher Scientific (see Table 1 for primer sequences).

**Table 1.** Taqman Primers. Taqman primers for *Mmp13, Mmp9, Rankl, Opg, Rank*, and *M-csf* are listed along with their amplicon lengths.

| *Gene* | *TaqMan assay* | *Amplicon length* |
| --- | --- | --- |
| *Mmp13* | Mm00439491_m1 | 65 |
| *Mmp9* | Mm00442991_m1 | 76 |
| *Rankl* | Mm00441906_m1 | 66 |
| *Opg* | Mm01205928_m1 | 75 |
| *Rank* | Mm00437132_m1 | 65 |
| *M-csf* | Mm00432686_m1 | 70 |

Expression of *Col1□1, Mmp13, Mmp9, Rankl, Opg, Rank*, and *M-csf* were normalized to expression of the reference gene *β-actin*. Amplification efficiencies were monitored across samples to ensure they remained sufficiently equal. Fold changes were calculated using the delta-delta C(t) method (37). To assess relative fold changes between genotypes, *Tgfbr1*^*fl/fl*^*;Wnt1-Cre* mice were compared relative to control mice at E18.5. Four control and four *Tgfbr1*^*fl/fl*^*;Wnt1-Cre* embryos from the same litter were assessed.

### 2.6 The use of quail embryos

Fertilized eggs of Japanese quail (*Coturnix coturnix japonica*) were acquired from the Michigan State University Poultry Farm (Lansing, MI) and incubated at 37°C in a humidified chamber until they reached embryonic stages appropriate for injections and analyses. The Hamburger-Hamilton (HH) staging system was used to properly stage quail embryos (38, 39). For all procedures, we adhered to accepted practices for the humane treatment of avian embryos as described in S3.4.4 of the *AVMA Guidelines for the Euthanasia of Animals*; 2013 Edition (40).

### 2.7 Inhibition of TGFBR1 in quail embryos

Quail eggs were windowed at HH9.5 using surgical scissors and transparent tape. Ten microliters of the TGFBR1 inhibitor (iTGFBR1) SB431542 (Sigma, SB) was injected into the vitelline vein of quail at stage HH33, which by Carnegie staging is similar to an E15 mouse, and therefore after the E10-E14.5 apoptosis phenotype seen in the *Tgfbr1*^*fl/fl*^*;Wnt1-Cre* mouse embryos (38, 41). The concentration was determined following dose-response studies and published literature. Control embryos were treated with vehicle dimethylsulfoxide (DMSO). After injections, eggs were closed with tape and incubated until they were collected in 10% formalin at stage HH38. Five control vehicle-injected quail and four iTGFBR1-injected quail were assessed. All samples were injected and collected at the same time.

### 2.8 Whole-mount staining of calcified bone using alizarin red

To analyze skull and lower jaw bone morphology of control and iTGFBR1-treated quail embryos, HH38 embryos were collected in water, eviscerated, and fixed in 95% ethanol. Acetone was used to remove fat tissue. Five control and four iTGFBR1-treated quail embryos were then stained with alizarin red to identify calcified bone. Tissue was cleared in 0.5% KOH and 0.06% H_2_O_2_, then processed through a Glycerol series (25%, 50%, 75%, 100%) (4, 5, 17, 27, 42).

Alizarin red-stained control and iTGFBR1-treated quail embryos were imaged on a MZ95 microscope (Leica, Buffalo Grove, IL) with a DP72 camera using cellSens Entry software (Olympus, Center Valley, PA).

### 2.9 Measurement of jaw length in quail embryos

Lower jaw length of n=5 control and n=4 iTGFBR1-treated quail embryos was quantified using ImageJ (43). Lower jaw length was assessed using lateral micrographs by measuring from the proximal tip of the angular bone to the distal tip of the dentary bone. Upper jaw length was assessed using the same lateral micrographs by measuring from the tip of the nasal bone in the center of the maxilla to the distal tip of the premaxilla. Proportional lower jaw length is depicted as lower jaw to upper jaw ratio to normalize for skull size.

### 2.10 Statistics

Bone stereology and osteoclast measures assessed with Bioquant Osteo data are represented as median ± interquartile range with all points shown and compared by the Wilcoxon rank sum test. Bone volume and tissue mineral density measures from micro-CT scans show mean relative to controls ± standard deviation. Gene expression data are shown by mean fold change ± standard deviation with all data points shown. Quail jaw proportion data are represented as median ± interquartile range with all points shown. Micro-CT, gene expression, and quail jaw proportion data was assessed by an unpaired Student’s *t-*test.

## 3 Results

*Tgfbr1*^*fl/fl*^*;Wnt1-Cre* embryos at E18.5 had a severe micrognathia phenotype and lacked all condylar, coronoid, and angular processes consistent with the phenotype of these mice previously described by Dudas et al., 2006 and Zhao et al., 2008 (Figure 1A & 1B). In addition to the mandibular changes in *Tgfbr1*^*fl/fl*^*;Wnt1-Cre* embryos, the craniofacial defects of *Tgfbr1*^*fl/fl*^*;Wnt1-Cre* embryos were severe and perinatal lethal (data not shown, (17, 27)). Masson’s Trichrome staining of control *Tgfbr1*^*fl/+*^ and *Tgfbr1*^*fl/fl*^*;Wnt1-Cre* mandibles showed dramatic gross changes in mandibular anatomy at the molar, incisor, and ventral levels of *Tgfbr1*^*fl/fl*^*;Wnt1-Cre* embryos (Figure 1C & 1D). Surprisingly, closer histological analysis of the blue-stained mandibular bone did not reveal a stark difference in bone architecture between control and *Tgfbr1*^*fl/fl*^*;Wnt1-Cre* E18.5 embryos (Figure 1E & 1D). Further, bone quality and stereology analysis revealed *Tgfbr1*^*fl/fl*^*;Wnt1-Cre* embryos did not differ from control embryos in bone perimeter per bone area, trabecular-like width, trabecular-like number, or trabecular-like separation, which further suggests bone quality did not significantly differ between control and *Tgfbr1*^*fl/fl*^*;Wnt1-Cre* E18.5 embryos (Figure 1G-1J). Additionally, expression of Collagen, Type I, Alpha 1 (*Col1□1*) and matrix metalloproteinase (*Mmp13*), markers of bone deposition and activity, did not differ in the mandible between control and *Tgfbr1*^*fl/fl*^*;Wnt1-Cre* E18.5 embryos (Figure 1K & 1L). These data suggest minimal changes in mandibular bone quality in *Tgfbr1*^*fl/fl*^*;Wnt1-Cre* mandibles despite severe micrognathia and perinatal lethal craniofacial defects.

**Figure 1.**
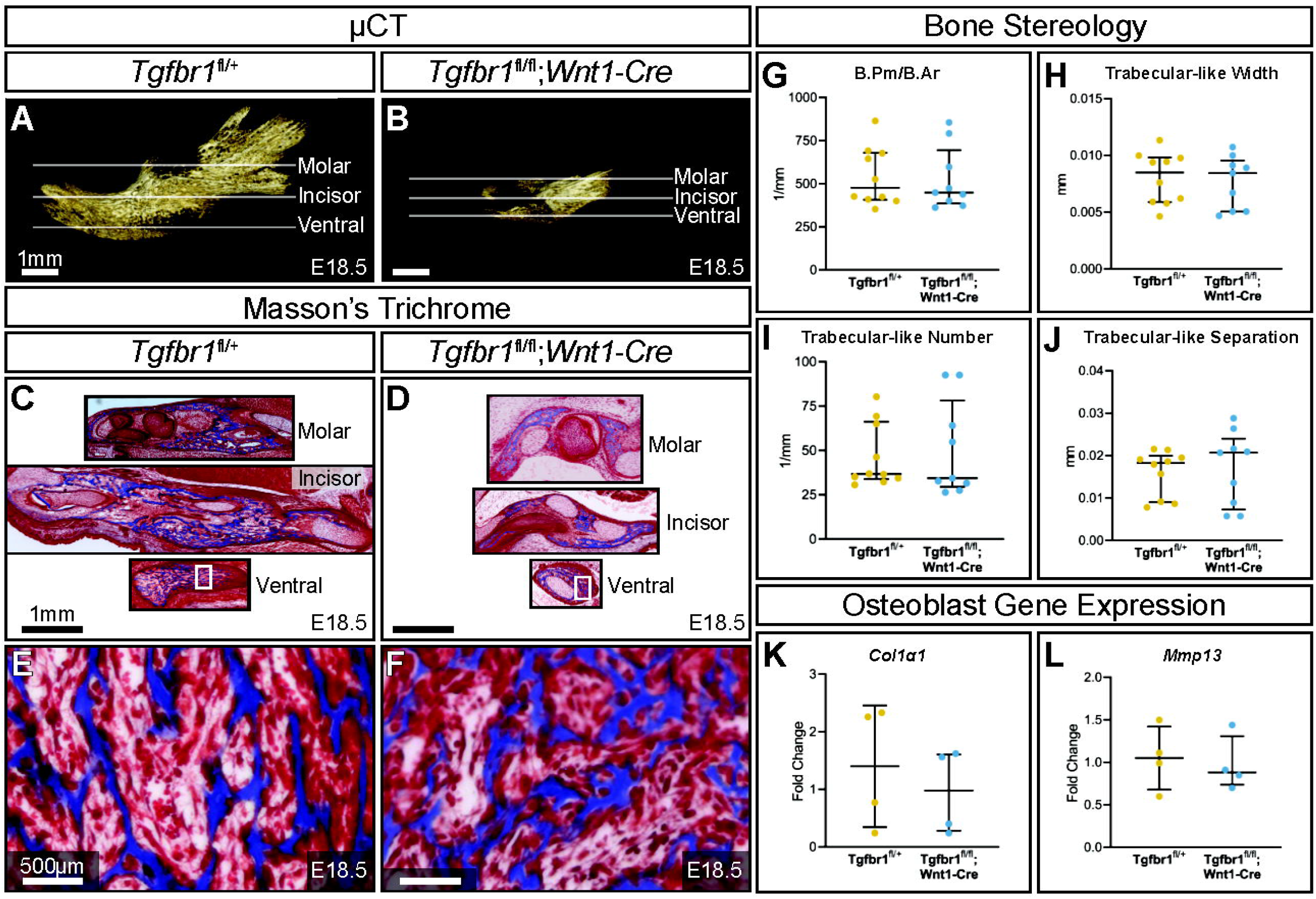
TGF-β signaling in neural crest does not affect mandibular bone quality. Lateral views of micro-CT scanned E18.5 (A) control and (B) *Tgfbr1*^*fl/fl*^*;Wnt1-Cre* mouse mandibles show differences in overall size and shape. Transverse sections were analyzed from three different regions in the mandible: molar, incisor, and ventral. Sections from the mandibles were stained with Masson’s Trichrome, which stains bone blue, from (C) control and (D) *Tgfbr1*^*fl/fl*^*;Wnt1-Cre* littermate E18.5 embryos. Higher magnification images from Masson’s Trichrome stained ventral sections (white boxes) are shown for the (E) control and (F) *Tgfbr1*^*fl/fl*^*;Wnt1-Cre* littermate E18.5 embryos. The sections were analyzed to determine if there were differences in bone stereology. There were no significant differences in (G) bone perimeter (B.Pm) per bone area (B.Ar), (H) trabecular-like width, (I) trabecular-like number, and (J) trabecular-like separation between the controls (blue) and *Tgfbr1*^*fl/fl*^*;Wnt1-Cre* mice (cKO, yellow). Using RT-qPCR to assay for (K) *Col1α1* and (L) *Mmp13* mRNA as a marker for bone deposition shows no significant difference in *Tgfbr1*^*fl/fl*^*;Wnt1-Cre* (yellow) at E18.5 compared to controls (blue).

To evaluate mandibular development between E16.5 and E18.5, embryonic control and *Tgfbr1*^*fl/fl*^*;Wnt1-Cre* mandibles were imaged by micro-CT to visualize the ossified bone at each of these stages (Figure 2A-2H). To evaluate mandibular bone resorption between E16.5 and E18.5, embryonic control and *Tgfbr1*^*fl/fl*^*;Wnt1-Cre* mandibles were stained for tartrate-resistant acid phosphatase (TRAP), which visualizes bone resorption in red (Figure 2I-2P). At both E16.5 and E18.5, the macroscopic spatial pattern of mandibular bone resorption did not greatly differ between embryonic control and *Tgfbr1*^*fl/fl*^*;Wnt1-Cre* mandibles despite drastic morphological differences in the mandible between the two genotypes. Bone stereology could not be compared between E16.5 and E18.5 because the E16.5 embryos lacked sufficient bone tissue for quantitative analysis. In addition to morphological differences in *Tgfbr1*^*fl/fl*^*;Wnt1-Cre* mandibles, bone volume was significantly decreased in *Tgfbr1*^*fl/fl*^*;Wnt1-Cre* mandibles at E16.5 and E18.5; *Tgfbr1*^*fl/fl*^*;Wnt1-Cre* mandibles had 86% less bone volume relative to controls at E16.5 (Figure 2Q) and 72% less bone volume relative to controls at E18.5 (Figure 2R). Though bone volume differed significantly between genotypes, tissue mineral density did not differ significantly between groups, which mirrors the lack of bone stereology differences between *Tgfbr1*^*fl/fl*^*;Wnt1-Cre* and control mandibles as well as the retained pattern of bone resorption in *Tgfbr1*^*fl/fl*^*;Wnt1-Cre* mandibles.

**Figure 2.**
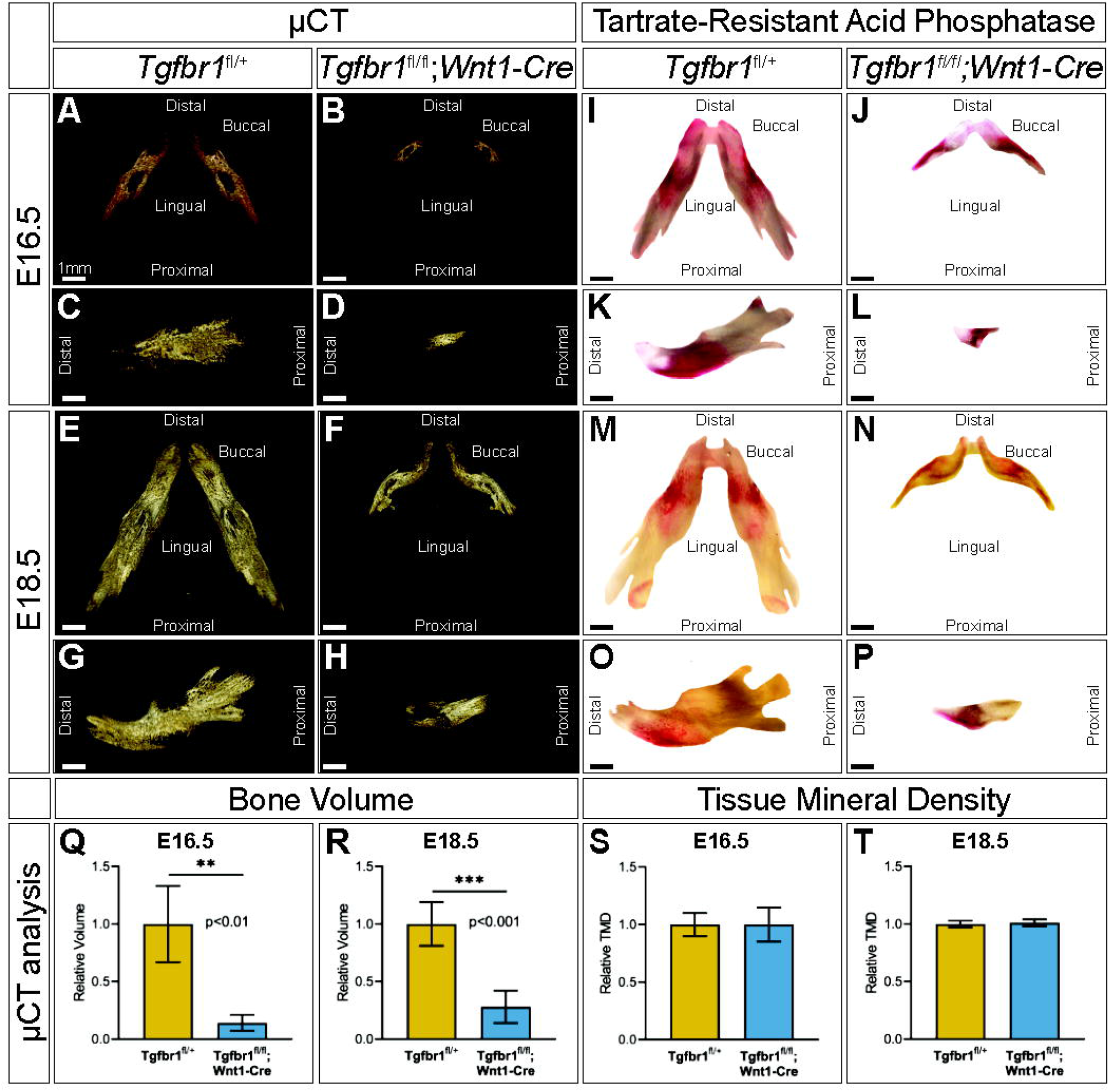
TGF-β signaling in neural crest does not control the spatial pattern of mandibular bone resorption or tissue mineral density. Micro-CT scanned mandibles from control and *Tgfbr1*^*fl/fl*^*;Wnt1-Cre* E16.5 littermate embryos, (A, B) dorsal and (C, D) lateral views, and E18.5 littermate embryos, (E, F) dorsal and (G, H) lateral views. Tartrate resistant acid phosphatase (TRAP) stained mandibles from control and *Tgfbr1*^*fl/fl*^*;Wnt1-Cre* E16.5 littermate embryos, (I, J) dorsal and (K, L) lateral views, and E18.5 littermate embryos, (M, N) dorsal and (O, P) lateral views. Micro-CT analysis of bone volume from (Q) E16.5 and (R) E18.5 control and *Tgfbr1*^*fl/fl*^*;Wnt1-Cre* embryos and tissue mineral density from (S) E16.5 and (T) E18.5 control and *Tgfbr1*^*fl/fl*^*;Wnt1-Cre* embryos.

TRAP-stained sections at the molar, incisor, and ventral level of the mandible (Figure 3A-3D) revealed on the microscopic level that the conditional knock out of *Tgfbr1* in NCM-derived osteoblasts resulted in a significant 3.2-fold decrease in osteoclast number per bone area in *Tgfbr1*^*fl/fl*^*;Wnt1-Cre* mandibles (Figure 3E). Additionally, osteoclast perimeter per bone area significantly decreased 3.2-fold in *Tgfbr1*^*fl/fl*^*;Wnt1-Cre* mandibles compared to controls (Figure 3F). These data show there are fewer osteoclasts within the same bone area in the *Tgfbr1*^*fl/fl*^*;Wnt1-Cre* mandibles. Receptor activator of nuclear factor Kappa-B (*Rank*) expression decreased 2.5-fold in *Tgfbr1*^*fl/fl*^*;Wnt1-Cre* mandibles, similarly suggesting a decrease in osteoclasts (Figure 3G). Yet matrix metalloproteinase 9 (*Mmp9*) expression decreased only 0.64-fold in *Tgfbr1*^*fl/fl*^*;Wnt1-Cre* mandibles, demonstrating a non-significant decrease in active bone resorption (Figure 3H).

**Figure 3.**
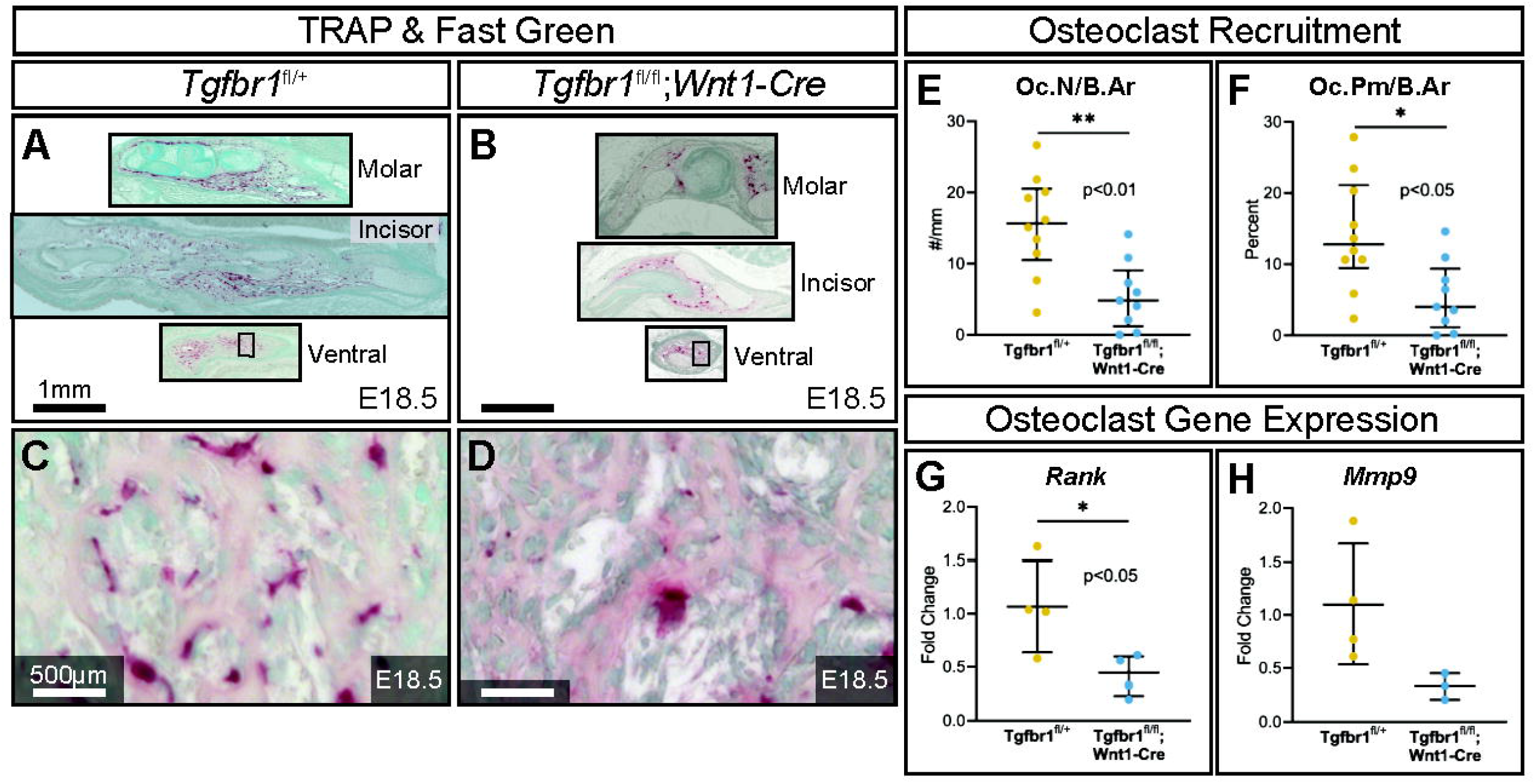
TGF-β signaling in neural crest controls the amount of mandibular bone resorption. Adjacent transverse sections from mandibles were that stained with Masson’s Trichrome were stained with Tartrate Resistant Acid Phosphatase (TRAP), which stains bone resorption red, and methyl green from (A) control and (B) *Tgfbr1*^*fl/fl*^*;Wnt1-Cre* E18.5 littermate embryos. Sections were stained and analyzed from three different regions in the mandible: molar, incisor, and ventral. Zoomed in views from TRAP and methyl green stained ventral sections (black boxes) are shown for the (C) control and (D) *Tgfbr1*^*fl/fl*^*;Wnt1-Cre* littermate E18.5 embryos. The sections were analyzed to determine if there were differences in the amount of bone resorption. There were significant differences in bone resorption, as analyzed by (E) osteoclast number per bone area (Oc.N/B.Ar) and (F) osteoclast perimeter per bone area (Oc.Pm/B.Ar), of an approximately 3.2-fold decrease for both in the *Tgfbr1*^*fl/fl*^*;Wnt1-Cre* mice when compared to control. (G) Using RT-qPCR to assay for *Rank* mRNA as a marker for number of osteoclasts shows a significant 2.5-fold decrease in *Tgfbr1*^*fl/fl*^*;Wnt1-Cre* at E18.5 when compared to controls. (H) Levels of *Mmp9* mRNA shows a 0.64-fold decrease in *Tgfbr1*^*fl/fl*^*;Wnt1-Cre* at E18.5 when compared to controls.

As osteoclasts are mesoderm-derived, not NCM-derived or targeted directly by Wnt1-Cre, they should retain *Tgfbr1*. Consequently, changes in osteoclasts in *Tgfbr1*^*fl/fl*^*;Wnt1-Cre* mandibles must be due to alterations in another cell type. Genes known to directly influence osteoclastogenesis were analyzed to investigate the possibility that changes in mesoderm-derived osteoclasts in *Tgfbr1*^*fl/fl*^*;Wnt1-Cre* mandibles were due to altered signaling from NCM-derived osteoblast lineage cells (“osteoblast-to-osteoclast signaling”). Expression of osteoblast- and osteocyte-secreted Rank ligand (*Rankl*) and *Rankl*’s inhibitor osteoprotegerin (*Opg*) did not differ significantly between *Tgfbr1*^*fl/fl*^*;Wnt1-Cre* and control mandibles (Figure 4A & 4B). As *Opg* inhibits *Rankl*, the *Rankl* to *Opg* ratio was also analyzed. Though the *Rankl* to *Opg* ratio trended downward in *Tgfbr1*^*fl/fl*^*;Wnt1-Cre* mandibles, the difference between *Tgfbr1*^*fl/fl*^*;Wnt1-Cre* and control mandibles was not significantly different (Figure 4C). Macrophage colony-stimulating factor (*M-csf*) similarly trended downwards in *Tgfbr1*^*fl/fl*^*;Wnt1-Cre* mandibles, but the decrease in *M-csf* expression in *Tgfbr1*^*fl/fl*^*;Wnt1-Cre* mandibles was not statistically significant (Figure 4D). Therefore, while the severely micrognathic *Tgfbr1*^*fl/fl*^*;Wnt1-Cre* mandibles had decreased osteoclast number per bone area, osteoclast perimeter per bone area, and *Rank* expression despite mesoderm-derived osteoclasts retaining *Tgfbr1*, the mechanism is not due to drastic changes in *Rankl, Opg, Rankl* to *Opg* ratio, or *M-csf* expression.

**Figure 4.**
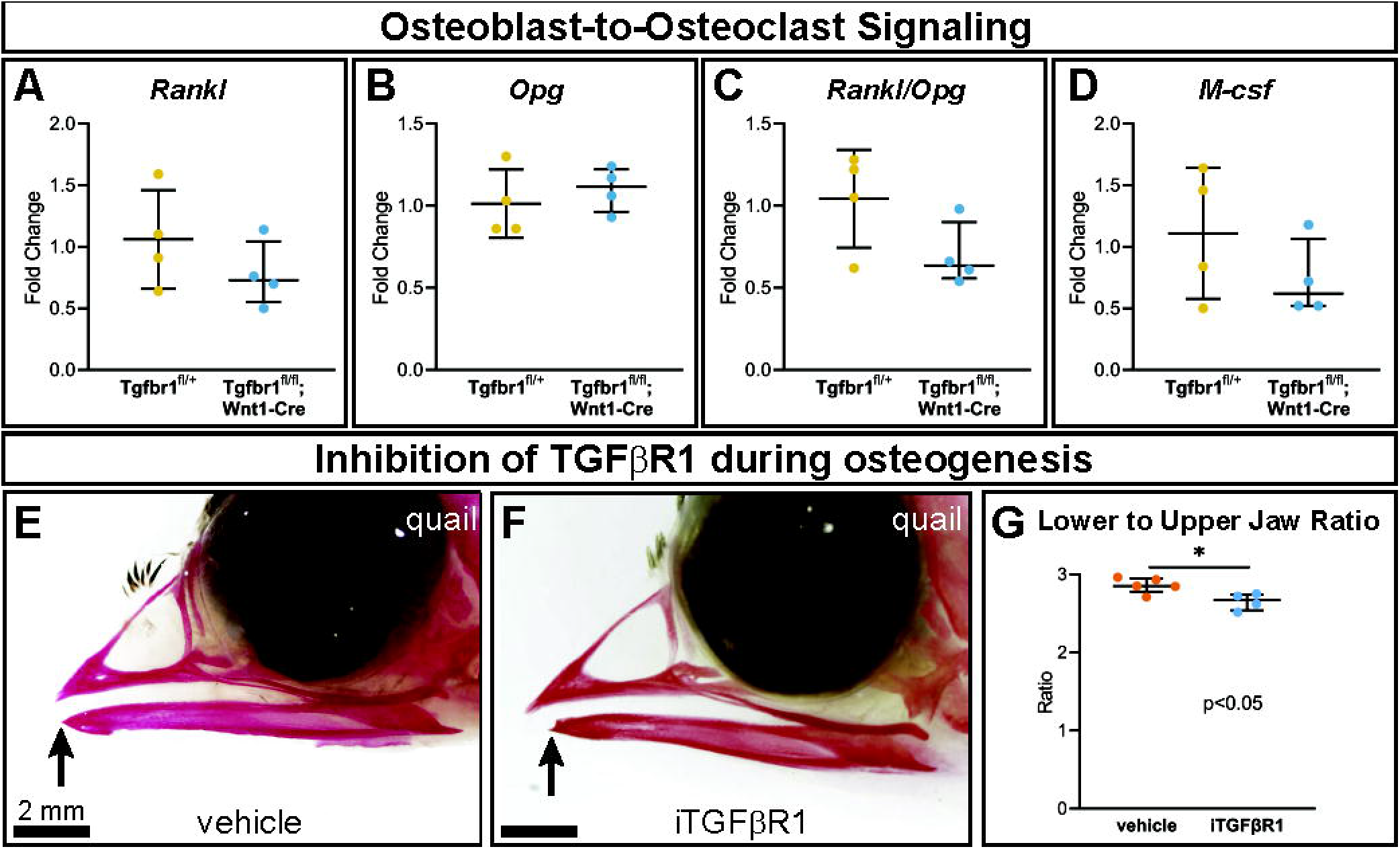
TGF-β signaling in neural crest does not alter osteoblast-to-osteoclast signaling pathways through *Rankl, Opg*, or *M-csf*. Well-known signaling factors from the osteoblast lineage to osteoclasts are Rankl, Opg, and M-csf. Using RT-qPCR to assay for (A) *Rankl*, (B) *Opg*, (C) *Rankl/Opg*, and (D) *M-csf* mRNA expression revealed no significant difference in *Tgfbr1*^*fl/fl*^*;Wnt1-Cre* at E18.5 compared to controls. To inhibit TGFBR1 after the apoptosis in *Tgfbr1*^*fl/fl*^*;Wnt1-Cre* mice from E10-E14.5, we injected a TGFBR1inhibitor (SB431542) to quail embryos at HH33, which by Carnegie staging is similar to an E15 mouse and therefore after the E10-E14.5 apoptosis. Embryos were collected at HH38 and stained with alizarin red, which stains calcified bone red. (E) In vehicle-treated control quail, the upper and lower jaws align, but in (F) TGFBR1inhibitor-injected quail the lower jaw is significantly shorter than the upper jaw. (G) The ratio of lower to upper jaw length is significantly less in the TGFBR1-inhibitor injected quail compared to the control injected quail.

To further interrogate the role of TGF-β signaling at a more specific developmental timepoint, a single dose of an inhibitor of TGFBR1 (iTGFBR1) or vehicle control was delivered to embryonic quail at the developmental timepoint HH33, which by Carnegie staging is similar to an E15 mouse. This timepoint is critically also after the E10-E14.5 apoptosis phenotype seen in the *Tgfbr1*^*fl/fl*^*;Wnt1-Cre* mouse embryos and just prior to osteoclast bone resorption beginning in the lower jaw. Several days after iTGFBR1 or vehicle treatment at developmental stage HH38 when the jaw is ossified, vehicle control-treated quail embryos stained with alizarin red demonstrate typical quail jaw morphology, with the distal tip of the lower jaw being aligned with the upper jaw (Figure 4E). In contrast, iTGFBR1-treated quail embryos have a shorter lower jaw compared to the upper jaw (Figure 4F). To account for potential changes in upper jaw length, jaw length was assessed as lower to upper jaw ratio. Quail embryos given a single treatment of iTGFBR1 had a significant, 6.3% decrease in lower to upper jaw ratio (Figure 4G). The decrease in iTGFBR1-treated quail lower jaw length recapitulates the micrognathia in *Tgfbr1* deficient *Tgfbr1*^*fl/fl*^*;Wnt1-Cre* mice embryos.

## 4 Discussion

It has previously been established in separate reports 1) that NCM-derived osteoblast lineage cells influence embryonic mandibular development by controlling osteoclastogenesis and bone resorption, and 2) that loss of TGFBR1 in NCM results in micrognathic jaws which lack the coronoid, condylar, and angular processes (5, 17, 27). In this study, we investigated the possibility that TGF-β signaling in NCM is a critical signaling pathway through which NCM-derived osteoblast lineage cells influence osteoclast recruitment and osteoclast activity during mandibular development. Despite drastic craniofacial and mandibular defects, *Tgfbr1*^*fl/fl*^*;Wnt1-Cre* mice which lacked the Tgfbr1 receptor in NCM had mandibular bone quality similar to controls, mandibular tissue mineral density similar to controls, and mandibular bone resorption which was macroscopically similarly located to bone resorption in control mandibles. However, closer histological investigation revealed *Tgfbr1*^*fl/fl*^*;Wnt1-Cre* mice had fewer osteoclasts per bone area, as well as significantly decreased expression of osteoclast-related gene *Rank*. Examination of NCM-derived factors which may be influencing osteoclastogenesis and bone resorption, such as *Rankl, Opg*, and *M-csf*, revealed no meaningful difference between *Tgfbr1*^*fl/fl*^*;Wnt1-Cre* mice and controls, which suggests other NCM-derived factors are mediating the change in osteoclast number and osteoclast perimeter in *Tgfbr1*^*fl/fl*^*;Wnt1-Cre* mice.

Our data suggest that the decrease in osteoclast number and osteoclast perimeter per bone area in *Tgfbr1*^*fl/fl*^*;Wnt1-Cre* mandibles detected at E18.5 might be the effort of the mandibular bone to maintain bone quality and density following the previous apoptosis at E10-E14.5 in the proximal and aboral regions of the *Tgfbr1*^*fl/fl*^*;Wnt1-Cre* mandibles (17, 27). Interestingly, though NCM-derived osteoblast lineage cells are known to regulate osteoclastogenesis, the lack of change in *Rankl, Opg, M-csf*, and *Rankl:Opg* in *Tgfbr1*^*fl/fl*^*;Wnt1-Cre* mandibles suggests osteoblast lineage sells may be doing so outside of modulating these factors. Wang et al. similarly found no change in expression of *Rankl* or *Opg* in *Tgfbr2*^*flfl*^*;Osx-Cre* alveolar bone despite a decrease in osteoclast number and a decrease in TRAP-indicated bone resorption in alveolar bone (44). Further, that *Tgfbr1*^*fl/fl*^*;Wnt1-Cre* mandibles have bone quality similar to control mandibles suggests the apoptosis noted by Dudas et al. and Zhao et al. may occur directly in the population of NCM destined to form the processes and part of the ramus portions of the developing mandible, while not affecting the morphogenic units from the alveolar and body of the mandible (17, 27, 33-35). Further research is required to explore the possibility that NCM-derived osteoblast lineage cells influence osteoclast differentiation and activity to compensate for apoptosis in the proximal and aboral regions of *Tgfbr1*^*fl/fl*^*;Wnt1-Cre* mandibles between stages E10 and E14.5. We observed no difference in mandibular tissue mineral density in *Tgfbr1*^*fl/fl*^*;Wnt1-Cre* embryos despite a decrease in osteoclast number, osteoclast perimeter per bone area, and *Rank* expression, which also merits further investigation into the mechanism through which osteoclast-deficient mandibles have tissue mineral density similar to that of controls. Additionally, unbiased techniques such as mRNA sequencing may indicate which factors are utilized by osteoblast lineage cells to influence osteoclasts at and before E18.5, the stage at which we observed a decrease in osteoclast number and perimeter.

Embryonic quail given a single dose of iTGFBR1, specifically at the developmental timepoint when the lower jaw is just beginning to mineralize and just prior to osteoclast bone resorption beginning in the lower jaw, had a short lower jaw. This demonstrates inhibition of typical TGFBR1 activity results in a shorter lower jaw despite the drug being delivered *after* the timepoint at which apoptosis was seen in the *Tgfbr1*^*fl/fl*^*;Wnt1-Cre* mandibles at E10-E14.5. Utilizing an inducible cre mouse model to excise *Tgfbr1* after E14.5 could reveal the role of NCM-controlled osteoclastogenesis and bone resorption in mandible development independent of the apoptosis seen in the proximal and aboral regions of *Tgfbr1*^*fl/fl*^*;Wnt1-Cre* mandibles at E10-E14.5 in a mammalian model. Such a model would provide more insight into whether TGF-β signaling in NCM is a critical pathway by which osteocytes influence osteoclast activity and therefore jaw length, as well as at which timepoints TGF-β signaling meaningly influences jaw development and length.

In summary, our experiment recapitulated the mandibular defects previously described in *Tgfbr1*^*fl/fl*^*;Wnt1-Cre* mice and explored the status of osteoclasts in the mandibles of *Tgfbr1*^*fl/fl*^*;Wnt1-Cre* mice. Though a decrease in osteoclast number, osteoclast perimeter, and *Rank* and *Mmp9* expression were detected in *Tgfbr1*^*fl/fl*^*;Wnt1-Cre* mandibles, NCM-derived osteoblast lineage signaling to osteoclasts was not indicated by a change in *Rankl, Opg*, or *m-Csf* expression. Further investigation would elucidate the mechanism by which NCM-derived osteoblast lineage cells influence osteoclasts during mandibular development when TGFBR1 signaling is altered. In addition, further investigation of TGFBR1 signaling will improve our understanding of Loeys-Dietz Syndrome I and improve our ability to treat patients with the syndrome. Growing evidence points towards osteoclasts having a role in determination of jaw bone length under the direction of NCM-derived osteoblast lineage cells, making additional research into the mechanisms by which this relationship occurs critical for understanding mandibular development, and in turn, for developing therapies for all individuals with mandibular malformations.

## Conflict of Interest

*The authors declare that the research was conducted in the absence of any commercial or financial relationships that could be construed as a potential conflict of interest*.

## Author Contributions

CJH, SG, and EEB compiled the data, performed the statistical analysis, and co-wrote the manuscript. CJH, SG, VK, and EEB analyzed the data. SG, VK, and EEB designed the experiments. SG and EEB performed the experiments. VK and EEB designed the study. EEB conceived and coordinated the study. All authors contributed to manuscript revision, read, and approved the submitted version.

## Funding

Funded in part by the University of Michigan Undergraduate Research Opportunity Program Biomedical and Life Sciences Sumer Research Opportunity Program to SG, NIDCR R01 DE13085 to VK, NICDR K08 DE021705 to EEB, and NIH/NCRR S10RR026475-01 to the University of Michigan School of Dentistry Micro-CT core.

## Acknowledgments

We thank J. Lane, S. Rajderkar, and P. Thomas for suggestions and technical support, B. Pierchala for use and technical support of the Zeiss Microscope and AxioVision software, C. Merceron and E. Schipani for use and technical support of the Bioquant Osteo software, B. Allen for use of egg incubators and microinjector, M. Lynch in the University of Michigan School of Dentistry Micro-CT Core for scans, C. Strayhorn in the Histology Core for Masson’s Trichrome staining, and D. Folk and S. Hall for writing support through the BioKansas Scientific Writing Program.

## Data Availability Statement

Datasets are available on request: The raw data supporting the conclusions of this article will be made available by the authors, without undue reservation.

## Notes

### Competing Interest Statement

The authors have declared no competing interest.

